# quantms-rescoring enables deep proteome coverage across protein quantification, immunopeptidomics, and post-translational modifications experiments

**DOI:** 10.64898/2026.01.12.698877

**Authors:** Chengxin Dai, Ralf Gabriels, Robbin Bouwmeester, Asier Larrea, Jonas Scheid, Henry Webel, Fuchu He, Lennart Martens, Oliver Kohlbacher, Mingze Bai, Linhai Xie, Timo Sachsenberg, Yasset Perez-Riverol

**Affiliations:** State Key Laboratory of Medical Proteomics, Beijing Proteome Research Center, National Center for Protein Sciences (Beijing), Beijing Institute of Lifeomics, 102206, Beijing, China; International Academy of Phronesis Medicine (Guangdong), 510320, Guangdong, China; CompOmics, VIB Center for Medical Biotechnology, VIB, Ghent, 9052, Belgium; Department of Biomolecular Medicine, Faculty of Medicine and Health Sciences, Ghent University, Ghent, 9052, Belgium; European Molecular Biology Laboratory, European Bioinformatics Institute, Wellcome Genome Campus, Cambridge, United Kingdom; Department of Biochemistry and Molecular Biology. Faculty of Science and Technology. University of the Basque Country (UPV/EHU), Bilbao, Spain; Department of Peptide-based Immunotherapy, Institute of Immunology, University and University Hospital Tübingen, Tübingen, Germany; Cluster of Excellence iFIT (EXC2180) Image-Guided and Functionally Instructed Tumor Therapies, University of Tübingen, Tübingen, Germany; Quantitative Biology Center (QBiC), University of Tübingen, Tübingen, Germany; Institute for Bioinformatics and Medical Informatics (IBMI), University of Tübingen, Tübingen, Germany; The Novo Nordisk Foundation Center for Biosustainability, Technical University of Denmark, Lyngby, Denmark; BioOrganic Mass Spectrometry Laboratory (LSMBO), IPHC UMR 7178, University of Strasbourg, CNRS, Strasbourg, 67000, France; Infrastructure Nationale de Proteomique ProFI - FR2048, Strasbourg, 67087, France; Department of Computer Science, Applied Bioinformatics, University of Tübingen, Tübingen, Germany; Institute for Bioinformatics and Medical Informatics, University of Tübingen, 72076 Tübingen, Germany; Institute for Translational Bioinformatics, University Hospital Tübingen, 72074 Tübingen, Germany; Chongqing Key Laboratory of Big Data for Bio Intelligence, Chongqing University of Posts and Telecommunications, Chongqing, China

**Keywords:** Proteomics, Reanalysis, Rescoring, Workflow, Machine learning

## Abstract

The growing volume of public proteomics datasets and the advent of novel machine learning (ML)-based methods create unprecedented opportunities for discovery through large-scale reanalysis. However, traditional desktop tools are increasingly insufficient for processing and integrating data at this scale. To address this challenge, we present a novel package, quantms-rescoring, that extends the cloud-native quantms workflow with a machine learning-based rescoring module. Unlike prior tools that rescore single-engine outputs, quantms-rescoring seamlessly integrates multiple search engines (SAGE, COMET, and MSGF+), performs automatic model selection, model fine-tuning, and scales reproducibly on cloud infrastructures. In quantms-rescoring, we rely on multiple fragment-ion intensity (AlphaPeptDeep and MS2PIP) and retention-time prediction (DeepLC) methods to improve results from multiple peptide database search engines. It features automatic model selection, fine-tuning, and retraining for MS/MS intensity and retention time prediction to select the best model for a given dataset. We applied the novel workflow to five representative datasets spanning DDA label-free quantification, TMT 10-plex isobaric labelling of tumor proteomics data, immunopeptidomics, phospho-proteomics, and unseen lysine malonylation experiments. We achieved a 16-22.8% increase in identified spectra, along with the quantification of 2191 additional phosphorylated peptides and 1337 phosphosites. In the tandem mass tag (TMT)-labeled clear cell renal cell carcinoma dataset, 76 novel differentially expressed multiple search engines identified proteins with quantms-rescoring. Additionally, novel 11,688 HLA-II potential binders were detected in the immunopeptidomics dataset by multiple search engines with quantms-rescoring. For unseen malonylation data, we reported more than 58.8% malonylation PSMs and 30.5% modification sites than COMET alone. Together, these results show that integrating multi-engine searches with machine learning-derived features can be combined in a scalable workflow that enhances identification, PTM localization, and quantification performance.

## Introduction

In recent years, the field of proteomics has experienced rapid growth in publicly accessible datasets and a shift toward studies analyzing larger sample cohorts. As of November 2025, over 64,000 datasets have been submitted to ProteomeXchange (PX) repositories, including an increasing number of large-scale submissions with more than 100 instrument files (1, 2). This acceleration exposes practical limitations of many widely used identification and quantification tools, which were designed for study-by-study analysis, are desktop-centric (3–5), lack containerized command-line interfaces, or are not readily orchestrated by cloud-native workflow systems such as Nextflow (6). Integration is further hindered by licensing and openness: many widely used tools are commercial or closed-source, which constrains distribution and community-driven maintenance (7). Although some vendors now offer cloud deployments, these remain proprietary and are less amenable to workflow-native, reproducible use.

To address these challenges, we recently developed quantms (8), an open-source, cloud-native workflow for massively parallel quantitative proteomics reanalysis. quantms automatically distributes computations across multiple computing nodes via Nextflow and is fully based on standardized, open formats such as SDRF (Sample and Data Relationship Format) (9), mzML (10), and mzTab (11). quantms supports major data-dependent acquisition (DDA) workflows, including label-free quantification (DDA-LFQ), isobaric labeling methods such as TMT and iTRAQ (DDA-plex), and data-independent acquisition (DIA) methods, as well as downstream analysis protocols encompassing peptide and protein identification, immunopeptidomics, and post-translational modification analyses such as phosphoproteomics. Its reproducible execution environments are defined using Docker and Singularity (12), adhering strictly to the FAIR (Findability, Accessibility, Interoperability, and Reusability) principles (13).

With the adoption of machine learning (ML) and deep-learning (DL) in the field of proteomics, numerous models have been developed to infer physicochemical properties of peptides in LC-MS experiments; e.g., MS²PIP (14, 15), AlphaPeptDeep (16), Prosit (17), and DeepLC (18) for fragment ion intensities and retention time prediction. These models allow us to quantitatively assess the agreement between the predicted values and experimentally observed data. The resulting similarity metrics can be encoded as discriminative features and are used in rescoring approaches for identification, such as in the widely used Percolator algorithm (19, 20). For quantitative analysis, QuantSelect selects optimal fragments by systematically integrating various features (eg. XIC correlation) via self-supervised deep learning to improve quantification in DIA proteomics (21). In addition to the ML algorithms, different packages, including Oktoberfest (22), MS²Rescore (14, 23), or MSBooster (24) packages have been developed to facilitate the integration of these algorithms in MS analysis tools. The incorporation of these algorithms into already existing workflows for immunopeptidomics (25), proteogenomics (26), and quantitative proteomics analysis has proven to improve peptide proteomics identification in tools such as FragPipe (24) or MHCquant (25).

While the original quantms workflow relied solely on search-engine scores from Comet (27), MS-GF+ (28), and Sage (29) in combination with Percolator (19), it did not exploit peptide properties such as retention time and fragment-ion intensity or spectrum features such as signal-to-noise ratio. Here, we present quantms-rescoring, a Python package and Nextflow module that integrates MS²PIP, AlphaPeptDeep, DeepLC, and MS²Rescore into the quantms multi-search-engine workflow. quantms-rescoring introduces automatic feature generation, model selection, and transfer learning, enabling scalable ML-based rescoring across diverse quantitative proteomics applications.

## Methods

### Benchmarking quantms-rescoring and quantms

We evaluate the performance of the quantms-rescoring on six publicly available benchmark datasets. Three datasets were obtained from the PRIDE Archive (1) and one from the CPTAC data portal. The PXD001819 (30) standard proteomics dataset was generated by spiking 48 Sigma UPS1 proteins into a yeast cell lysate background at nine defined concentrations (0.05, 0.125, 0.25, 0.5, 2.5, 5, 12.5, 25, and 50 fmol/µL). Because spike-in levels are known, it serves as a benchmark for statistical evaluation of label-free identification and quantification performance. PXD019643 (31) is a large-scale immunopeptidomics dataset of matched HLA-I and HLA-II ligandomes from benign tissues from the HLA Ligand Atlas. HLA ligandome data from 29 distinct tissues collected from 21 individuals, totaling 1,274 LC-MS/MS runs across 225 mostly paired HLA-I (198) and HLA-II (220) samples. For the evaluation, we only used the HLA-II data. The PXD026824 (32) dataset comprises phosphopeptide-enriched samples used to evaluate post-translational modifications (PTM) identification and localization performance. PDC000127 is a large-scale TMT10 plex-labeled clear cell renal cell carcinoma cohort from the CPTAC repository. It includes 230 samples analyzed in 575 LC-MS/MS runs and was used to assess workflow performance in isobaric labeling experiments and large-cohort reanalysis scenarios. PXD021013 (33) is a synthetic HLA-I and HLA-II peptide dataset from the ProteomeTools project, which serves as a benchmark for evaluating false discovery rate (FDR) quantile control (33). Similarly, PXD009449 is a synthetic phosphorylated peptide dataset from the same project, which can be used to assess both FDR quantile control and false localization rate (FLR) (34). The PXD015089 dataset comprises malonylated proteins derived from *Toxoplasma gondii*, enriched via affinity-based methods, and serves to evaluate unseen PTM identification and localization performance (35).

To assess the impact of different search engine combinations and rescoring strategies, we evaluated six distinct quantms workflow configurations (**Figure 1**): (1) Comet alone, (2) Comet and MSGF+ combined, (3) Comet with quantms-rescoring, (4) Comet and MSGF+ with quantms-rescoring, and (5) Comet, MSGF+, and Sage with quantms-rescoring (**Figure 1A**). The different configurations enabled us to assess the impact of combining results from multiple search engines and of applying quantms-rescoring, which integrates ML-based features to improve peptide and protein identification and quantification performance. The Sage search engine was excluded from the immunopeptidomics datasets because its high memory requirements made it unsuitable for large-scale non-tryptic peptide searches, which involve substantially larger search spaces. All database search parameters were kept consistent with those described in our previous publication. For LFQ, TMT, and phosphorylation datasets, a 1% FDR threshold was applied at both the peptide-spectrum match (PSM) and protein group levels. For phosphorylation and lysine malonylation experiments, FLR was estimated, and a 1% threshold was applied. For immunopeptidomics, a 1% FDR threshold was applied at the PSM and peptide level. For the synthetic HLA peptide experiment, quantms searches raw files against the synthetic peptides combined with the UniProt human database. For the synthetic phosphorylated peptide experiment, quantms searches the raw files against a combined synthetic peptide and UniProt human database. Phospho (Y) was set as a variable modification.

**Figure 1:**
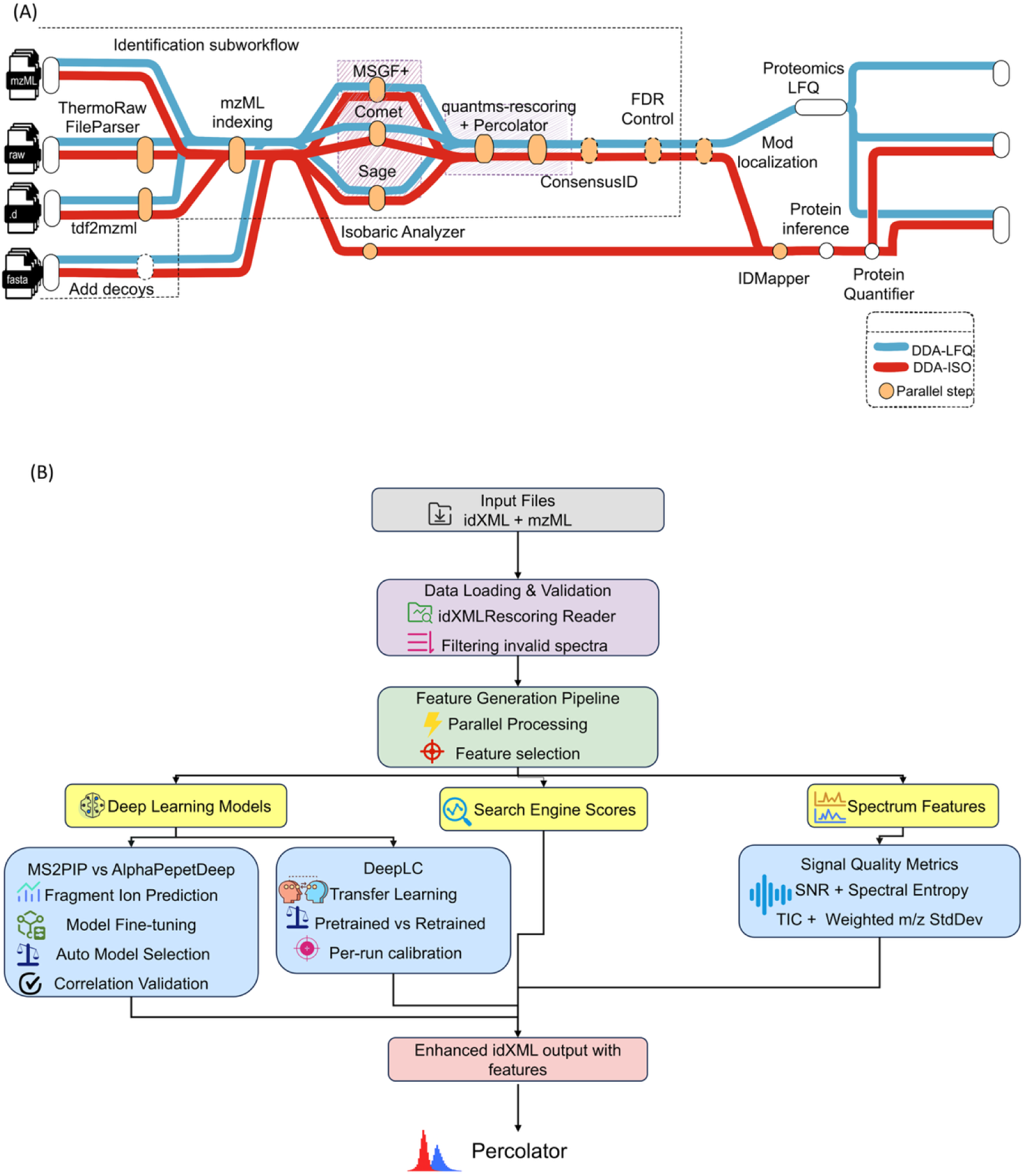
Overview of the quantms DDA workflow and rescoring module. (A) quantms DDA workflow includes three major steps: (1) peptide identification (using multiple search engines and quantms-rescoring with Percolator for boosting the number of identifications), (2) FDR control at PSM and peptide level, and (3) protein quantification for both TMT and label-free approaches. (B) The quantms-rescoring module loads search-engine input and results (idXML + mzML), validates them, and generates features in parallel. Features are derived from three sources: (1) deep-learning models - MS²PIP or AlphaPeptDeep for fragment-ion prediction with fine-tuning and DeepLC for retention-time transfer learning with automatic model selection and calibration; (2) search-engine scores; and (3) spectrum-quality metrics such as signal-to-noise ratio, spectral entropy, and others. The idXML files with additional features are then rescored by Percolator.

### Feature generation and model optimization in quantms-rescoring

quantms-rescoring calculates a comprehensive set of features based on the deviation between predicted and observed values, including retention times and fragment ion intensities (**Figure 1B**). In addition, the library allows the user to calculate and integrate auxiliary spectrum features (36) like signal-to-noise ratio (SNR) and spectrum entropy, which can be beneficial for specialized applications like immunopeptidomics or proteogenomics (**Supplementary Table 1**). quantms-rescoring introduces strategies for model selection, feature export, and model training to optimize prediction accuracy. For retention time (RT) modelling, quantms-rescoring uses transfer learning and retraining in DeepLC. It automatically evaluates available pre-trained models on a set of calibration PSMs, typically the top 20% scored PSMs, selects the model with the lowest mean absolute error (MAE), and then fine-tunes it on the current dataset to adapt to experiment-specific chromatographic conditions.

Unlike RT modeling, in fragment ion prediction (using tools such as AlphaPeptDeep and MS²PIP), quantms-rescoring selects the most suitable model for each MS run parameter. This approach is particularly important during reanalysis, as experimental metadata are often missing or incorrectly annotated - for instance, the fragmentation mode (e.g., HCD or CID) - or, in the case of very large experiments, analytical and instrument configurations may be mixed. quantms-rescoring selection algorithm takes the top 20% of peptide identifications and, with the model provided by the user, predicts the fragment-ions; then the Pearson correlation between the experimental and predicted; the model is rejected if at least 80% (default parameter) of the peptides do not correlate higher than a certain cutoff (default 0.6). When a model is rejected, the algorithm moves to the next one in MS^2^PIP or AlphaPeptDeep. In addition, quantms-rescoring seamlessly supports model fine-tuning to generate a specific-project model. This capability is crucial for applications such as post-translational modification (PTM) analysis, because existing MS/MS prediction models primarily cover common modifications, while the fragmentation patterns of modified peptides can diverge from those of unmodified peptides to varying extents depending on the modification type.

quantms-rescoring samples MS/MS spectra randomly from the full dataset based on a user-specified number of MS runs and partitions the sampled data into training and test sets according to a predefined ratio (default 0.6 for train) when enabling fine-tune. The training dataset is also provided by the user. Pre-trained and fine-tuned models are then evaluated and compared on the test set, after which the optimal model parameters are selected and retained. If no model meets this validation criterion when applying models, intensity-based rescoring is skipped unless explicitly enforced by the user, thereby preventing the application of poorly fitting or incompatible pre-trained models. This safeguard prevents the use of models that are poorly matched to the underlying data; for example, applying a CID model to an HCD MS run or using a model trained on human data for a distantly related species dataset, which could otherwise reduce identification rates or low-quality peptide identifications. compromise scoring accuracy. This is particularly important for large datasets that employ multiple fragmentation modes, PTMs, or analytical methods. All feature generation and model-selection steps are fully parallelized at the MS-run level, significantly reducing runtime for large-scale datasets. After feature extraction, the combined feature tables are passed to Percolator, which estimates FDR and posterior error probabilities (PEP).

During benchmarking, different workflow configurations were tested: search engines + Percolator only (no features from quantms-rescoring); search engines + quantms-rescoring + Percolator; and the last combination with additional spectrum SNR features. To merge results from multiple search engines, ConsensusID aggregates PSMs into unified scores based on the calculation of PEP and chooses the PSM with the highest probability. For phosphoproteomics datasets, LuciPHOr2 (37) is employed to assign site-FLR using OpenMS tools. All runtime parameters of quantms for different datasets are summarized in **Supplementary File 2**.

## Results

### quantms-rescoring enhances LFQ identification and quantification performance

To evaluate the performance of the quantms-rescoring-enhanced workflow at both identification and quantification levels, we analyzed the public UPS1 protein spike-in dataset PXD001819 (30). For benchmarking PSM identification performance, six workflow configurations that employ Percolator during post-scoring were compared: (1) Comet alone, (2) Comet and MSGF+ combined, (3) Comet with quantms-rescoring, (4) Comet and MSGF+ with quantms-rescoring, (5) Comet, MSGF+, and Sage and (6) Comet, MSGF+, and Sage with quantms-rescoring features. These configurations were tested to assess which fragment intensity-based and retention time-based features improve identification sensitivity and specificity. To compare workflow configurations, PSMs were filtered at different FDR thresholds. As shown in **Figure 2A**, using consensus scores (38) from multiple search engines increased the number of identifications at a fixed 0.01 FDR threshold (**Supplementary Figure 1**). Specifically, combining Comet, MSGF+, and Sage resulted in a 17.9% increase in identified spectra relative to Comet alone. Incorporating MS²PIP and DeepLC features through quantms-rescoring provided an additional 10.7% increase. Overall, quantms achieved 28.6% more PSM identifications than MaxQuant, demonstrating the benefits of combining multi-engine search with machine learning-based rescoring.

**Figure 2:**
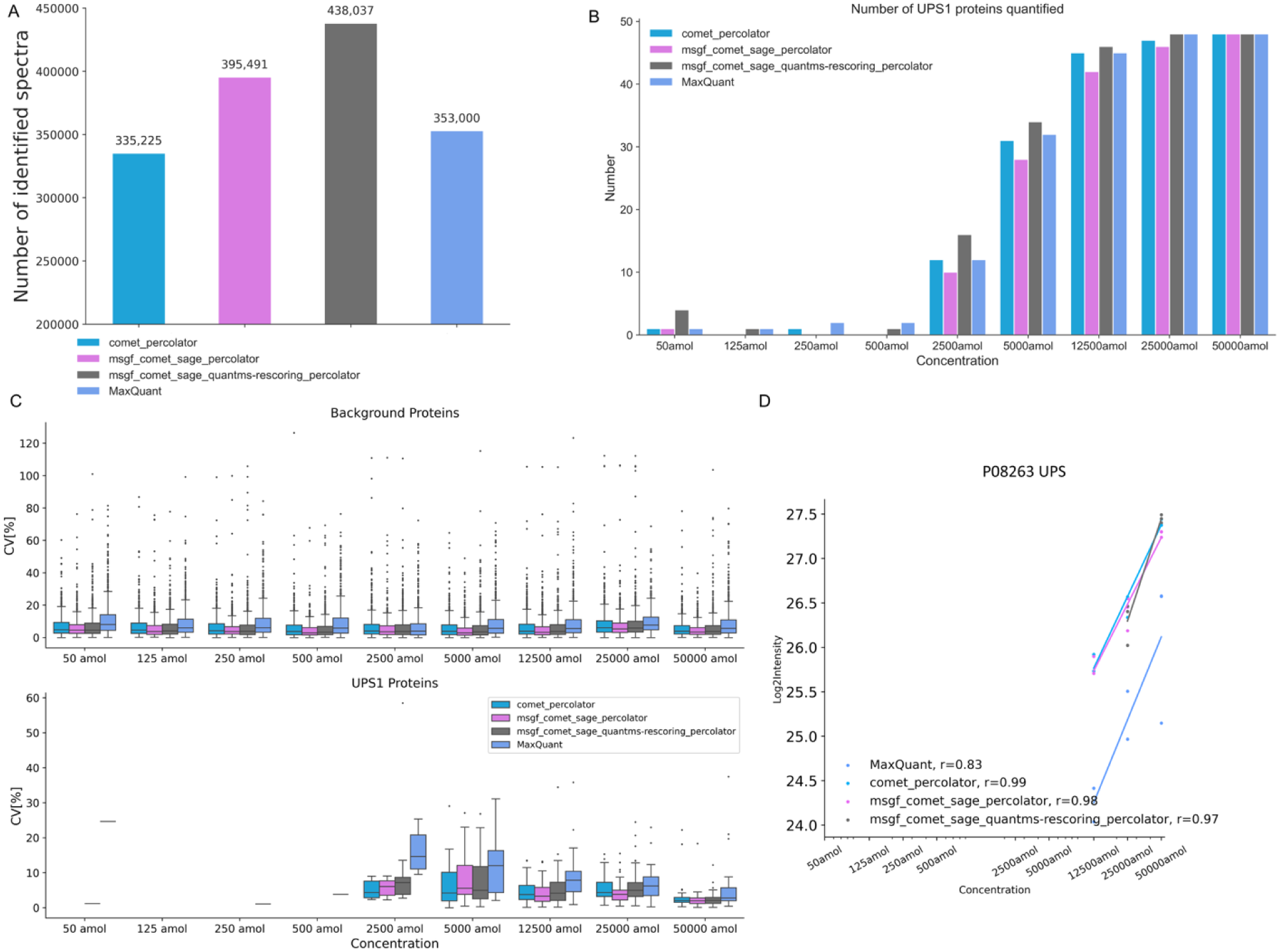
Comparison of identification and quantification results for different workflow settings on the PXD001819. (A) The number of identified spectra at 0.01 FDR levels for different workflow settings. (B) Protein quantification results for the PXD001819 dataset. Three different workflow settings are shown: Comet alone, three search engines, and the combination of three search engines with quantms-rescoring. All using Percolator in the rescoring process (C) Coefficient of variation of Yeast and UPS1 proteins. Only a very small number of data points are observed due to few UPS1 proteins are quantified when concentrations were below 2500 amol/µL. (D) The correlation between theoretical abundance and calculated protein abundance at different concentrations for the reanalysis of PXD001819 using MaxQuant and quantms with different configurations.

When benchmarking quantification results, the configurations that included quantms-rescoring-derived features enabled the detection and quantification of a greater number of low-abundance UPS1 proteins (**Figure 2B**). For example, multiple search engines with quantms-rescoring, compared to the settings without quantms-rescoring, and MaxQuant, can quantify up to 6 UPS1 proteins in 2500amol /µL and 4 more UPS1 proteins in 2500amol/µL, respectively. Only a small number of UPS proteins were quantified when the spike-in concentrations were below 2500 amol/µL, consistent with observations reported in the original research (30). The coefficients of variation (CVs) don’t change significantly before or after enabling quantms-rescoring, and quantms exhibited lower CVs compared to MaxQuant when considering only proteins quantified in at least 50% of replicates (**Figure 2C**). The correlation between theoretical abundance and calculated protein abundance at different concentrations doesn’t change significantly before or after enabling quantms-rescoring, and its performance was comparable to MaxQuant (**Figure 2D**). Overall, enabling quantms-rescoring did not cause a significant overall change in the variability of quantified proteins, suggesting that the rescoring process enhances identification confidence without introducing additional quantification bias.

To better understand the contribution of individual features to rescoring performance, we examined the top 20 feature weights from Percolator (**Supplementary Figure 2**). More than half of these features originated from MS^2^PIP and DeepLC, reflecting the strong influence of fragment intensity- and retention time-based predictors on model discrimination. Notably, *SpecPearsonNorm* had a strong positive weight, indicating that higher correlations between predicted and observed fragment intensities increase confidence in true PSMs. Conversely, *RtDiffBest* had a negative weight, suggesting that large retention time deviations reduce match reliability.

### Improved identification and quantification in TMT experiments via quantms-rescoring

We next applied the workflow to the large-scale TMT-labeled dataset from CPTAC (PDC000127). The integration of multiple search engines (Comet, MSGF+, Sage) and quantms-rescoring led to substantial improvements in both identification and quantification (**Figure 3**). Compared to Comet alone, integrating multiple search engines increased the PSM identification rate by 9%, resulting in 825 additional quantified proteins. The PSM identification rate increased by 5%, and 544 additional proteins were quantified compared to Comet, MSGF+, and Sage workflow configuration when further enabling quantms-rescoring (**Figure 3A-B, Supplementary Figure 3**). We then compared the quantified protein abundances obtained using the Comet, MSGF+, and Sage with and without quantms-rescoring methods (**Figure 3C**). As shown in the figure, six samples were randomly selected for display. Among these proteins quantified only by workflow with quantms-rescoring, an average of 103 proteins had abundance levels within the top 20% of all proteins for each sample (**Figure 3C**). We next assessed whether these improvements have the potential to translate into meaningful biological insights by conducting differential expression on the resulting quantification data. As shown in **Figure 3D**, Comet, MSGF with a quantms-rescoring workflow add 76 differentially expressed proteins (adj. p-value < 0.05 and log2FC > 1) compared to Comet, MSGF+ configuration, such as FOXG1, TLR10, and GABRB3 proteins.

**Figure 3:**
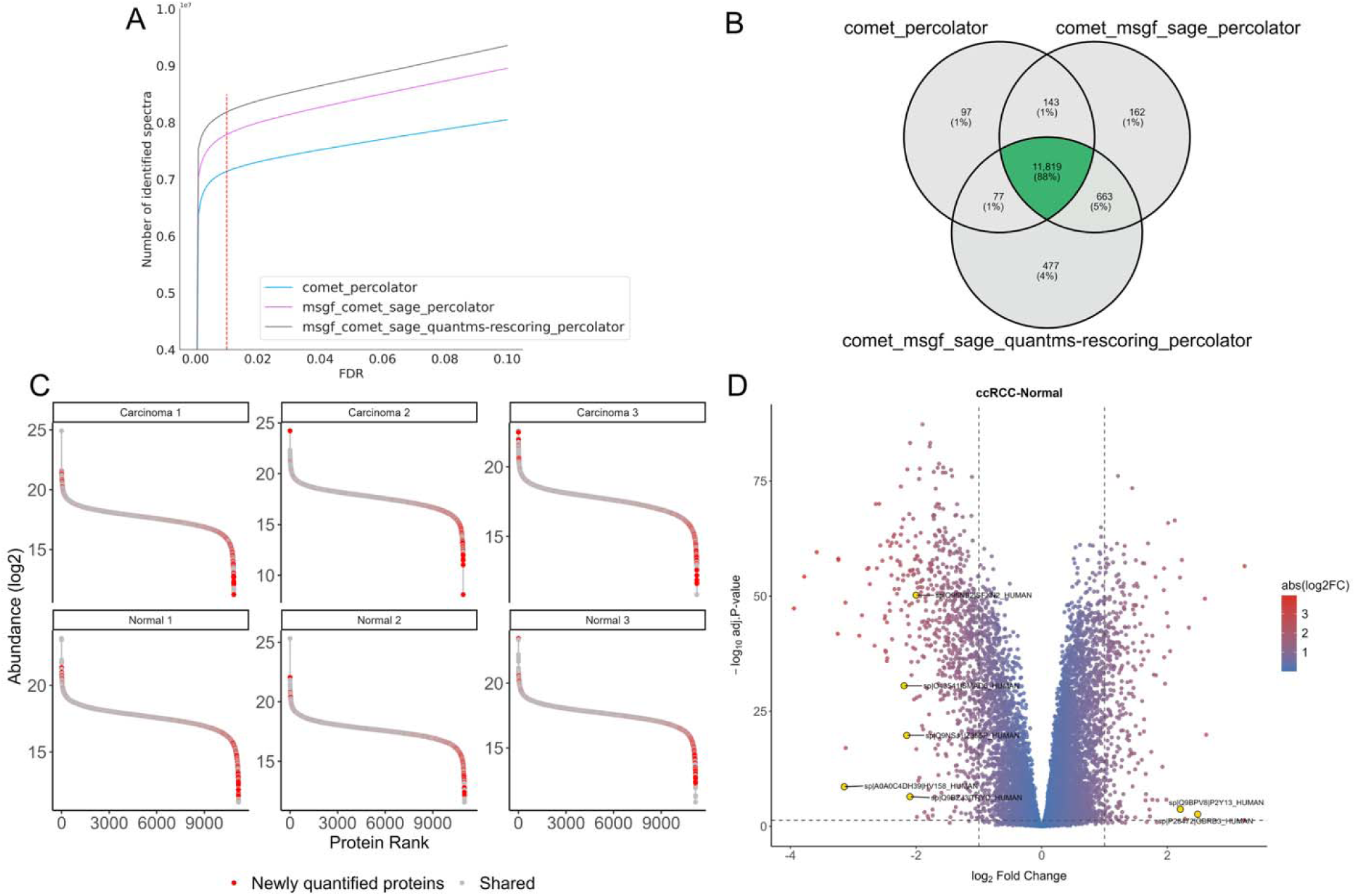
Comparison of identification and quantification results for different workflow settings on the CPTAC dataset PDC000127. (A) Number of identified spectra at different FDR thresholds and workflow settings. (B) The Venn plot of quantified proteins for three workflow settings. (C) Protein abundance rank comparison by using Comet, MSGF+, and Sage with and without quantms-rescoring for six samples. Red dots represent proteins quantified only in the workflow with quantms-rescoring. (D) Volcano plot generated with MSstatsTMT. Yellow dots denote representative differentially expressed proteins identified exclusively in the workflow incorporating quantms-rescoring. Differentially expressed proteins are defined as those with an adjusted p-value < 0.05 and log2FC > 1.

Notably, FOXG1and GABRB3 have been reported to be associated with prognosis in clear cell renal cell carcinoma (39, 40). TLR10 has been reported to be a biomarker for early diagnosis and prognosis assessment of renal cell carcinoma (41). These findings demonstrate that quantms incorporating quantms-rescoring enhances not only identification sensitivity but also has the potential to uncover proteins of clinical relevance. Feature importance (**Supplementary Figure 4**) revealed similar patterns as for the LFQ dataset, with strong weights on fragment intensity and retention time predicted features across multiple search engines.

### Multi-Engine consensus rescoring with quantms enhances peptide identification in HLA Class II Immunopeptidomics

The HLA Ligand Atlas (31) (PXD019643) provides a comprehensive resource of HLA-I and HLA-II presented peptides derived from 30 benign tissues, 51 HLA-I alleles, and 86 HLA-II alleles. To evaluate the performance of the quantms workflow with quantms-rescoring, we analyzed the HLA Class II subset of the HLA Ligand Atlas. Five workflow configurations were evaluated, all using Percolator in the rescoring process: (1) Comet alone, (2) Comet with quantms-rescoring features, (3) Comet and MSGF+, (4) Comet and MSGF+ with quantms-rescoring features, and (5) Comet and MSGF+ with quantms-rescoring and SNR features. Performance was assessed by the total number of PSMs and the number of unique peptide sequences identified. Across all comparisons, combining multiple search engines with quantms-rescoring features consistently improved identification performance. At 1% local PSM FDR thresholds, integrating MSGF+ with Comet increased identified PSMs by 11.7% relative to using Comet alone. Adding quantms-rescoring features to the multi-engine workflow yielded an additional 22.8% improvement (see **Figure 4A**).

**Figure 4:**
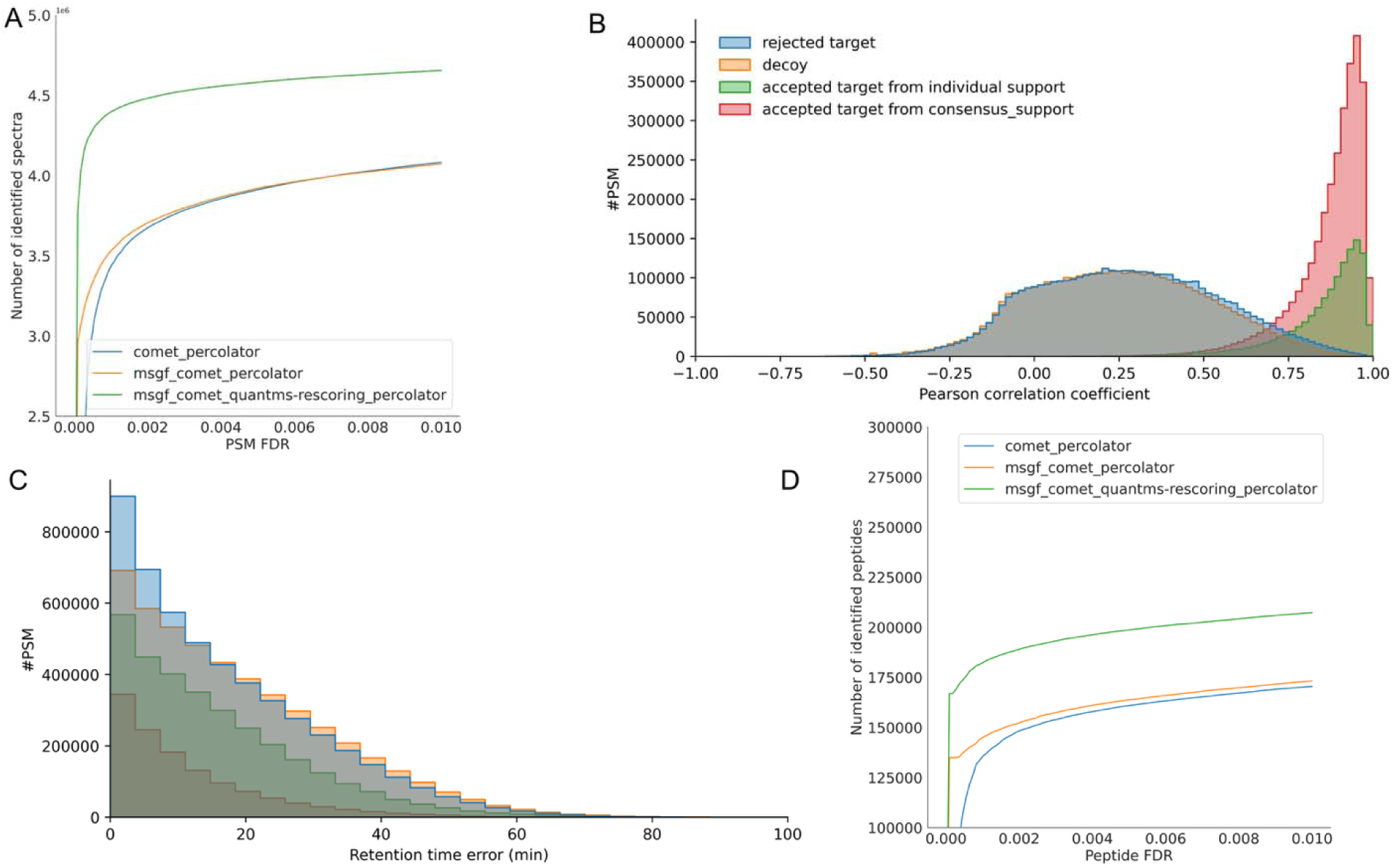
Comparison of immunopeptide identification results for different workflow settings on PXD019643. (A) Number of PSMs identified at different PSM FDR thresholds. (B) Pearson correlation between observed and predicted peak intensities. (C) Distribution of the smallest relative retention time error between observed and predicted retention time. Decoys (orange) rejected targets (blue; q-value *>*0.01), accepted targets (green: single search engine; red: multiple search; q-value *<=* 0.01). (D) Number of peptides identified at different peptide FDR thresholds.

Furthermore, we evaluated and compared FDR quality control for different workflow configurations on PXD021013 synthetic HLA-I and HLA-II peptide datasets (33). The results are shown in Supplementary Table 2. The estimated FDR (1.2% at the PSM level and 1.7% at the peptide level) is very close to the actual FDP when the FDR threshold is set to 1%, and the introduction of multiple search engines and rescoring does not disrupt the control of FDP. At 1% PSM FDR thresholds and 1% peptide level FDR, combining MS-GF+ and Comet with quantms-rescoring yielded 18.4% and 16.5% more identified PSMs and peptides, respectively, than using Comet alone.

To elucidate how quantms-rescoring features contribute to performance improvements, we examined the distributions of decoy, rejected target, and accepted target PSMs using two key metrics: the Pearson correlation coefficient (PCC) between predicted and observed fragment intensities, and the relative retention time error between predicted and observed retention times (**Figure 4B, C**). Accepted target PSMs were clearly distinguished from both decoy and rejected targets based solely on PCC values (median PCC 0.90 in accepted target PSMs, 0.2 in rejected target PSMs and 0.2 in decoy PSMs) and consistently displayed minimal retention time deviations (median retention time error 11 min in accepted target peptides, 14 mins in reject peptides, and 16 mins in decoy peptides), demonstrating the model’s effective discriminative capability. The top 20 feature weights were shown in Supplementary **Figure 5** from Percolator, revealing strong contributions of MSGF+ features such as *SpecEValue*, and suggesting that there are limited gains when MSGF+ is used alone. In contrast, nine MS2PIP-derived and two DeepLC-derived features also dominated the weight distribution, such as *RtDiffBest* and *DotProdIonBNorm* when Comet is used alone, demonstrating that consensus rescoring effectively leverages complementary information across search engines.

Furthermore, with a local peptide-level FDR threshold of 1% among the technical replicates per sample, integrating MSGF+ with Comet increased the number of identified peptides by 1.6% compared to using Comet alone. Adding quantms-rescoring features on top of the Comet workflow yielded a further 14% improvement compared to MSGF+ with the Comet workflow (**Figure 4D**). Enabling the quantms-rescoring combined with MSGF+ and Comet, the peptide identification improved by an additional 5% compared to workflows using only Comet and quantms-rescoring. When considering spectra features such as SNR in **Supplementary Table 1**, the peptide identification improved by an additional 0.6% compared to workflows using MSGF+, Comet, and quantms-rescoring (**Supplementary Figure 6A**).

To demonstrate the application potential of the quantms pipeline for antigen discovery, we show the number of HLA class II peptides predicted as binders (% binding rank ≤5) by the epitope prediction pipeline (**Supplementary Figure 6B-D**) (34). NetMHCIIpan 4.3 (35) was specified as the prediction tool. Added quantms-rescoring features to the Comet workflow, enabling the identification of 8,867 additional binders. When quantms-rescoring was enabled in combination with MSGF+ and Comet, an additional 2,821 binders were identified. And the length distributions of HLA class II peptide binders predicted are consistent with different workflow configurations. It displays a local maximum occurring at approximately 15-mers (**Supplementary Figure 6E**). Collectively, these findings demonstrate the robustness of the quantms workflow for immunopeptidomics applications and highlight its strength in leveraging machine-learning-derived features through consensus rescoring to enhance the identification accuracy of potential binders.

### Performance of the quantms-rescoring workflow in phosphoproteomics

We further evaluated the performance of the quantms workflow with quantms-rescoring on a PTM dataset (PXD026824). The AlphaPeptDeep model is used when enabling quantms-rescoring. For phosphoproteomics analysis, four workflow configurations, all using Percolator in rescoring, were benchmarked: (1) Comet alone, (2) Comet with quantms-rescoring features, (3) Comet and MSGF, and (4) Comet and MSGF with quantms-rescoring features. As shown in **Figure 5 and Supplementary Figure 7**, combining multiple search engines substantially improved Percolator’s ability to distinguish true from false PSMs. This improvement is reflected in a leftward shift in score distributions relative to the Comet-only workflow (**Figure 5A**). Incorporating quantms-rescoring features into the two-search-engine consensus further increased the number of identified spectra by 24%. Phosphorylated peptides frequently exhibit a loss of phosphoric acid during fragmentation (-98 Da neutral losses). To assess its influence on rescoring, quantms-rescoring calculates features (*modloss)* that account for these ions. Including those results in a moderate increase in the number of identified spectra by 3%. Feature importance analysis (see **Supplementary Figure 8**) shows similar trends as for the other datasets, highlighting the consistent importance of features such as *IonYNorm* and *DotProdIonYNorm*.

**Figure 5:**
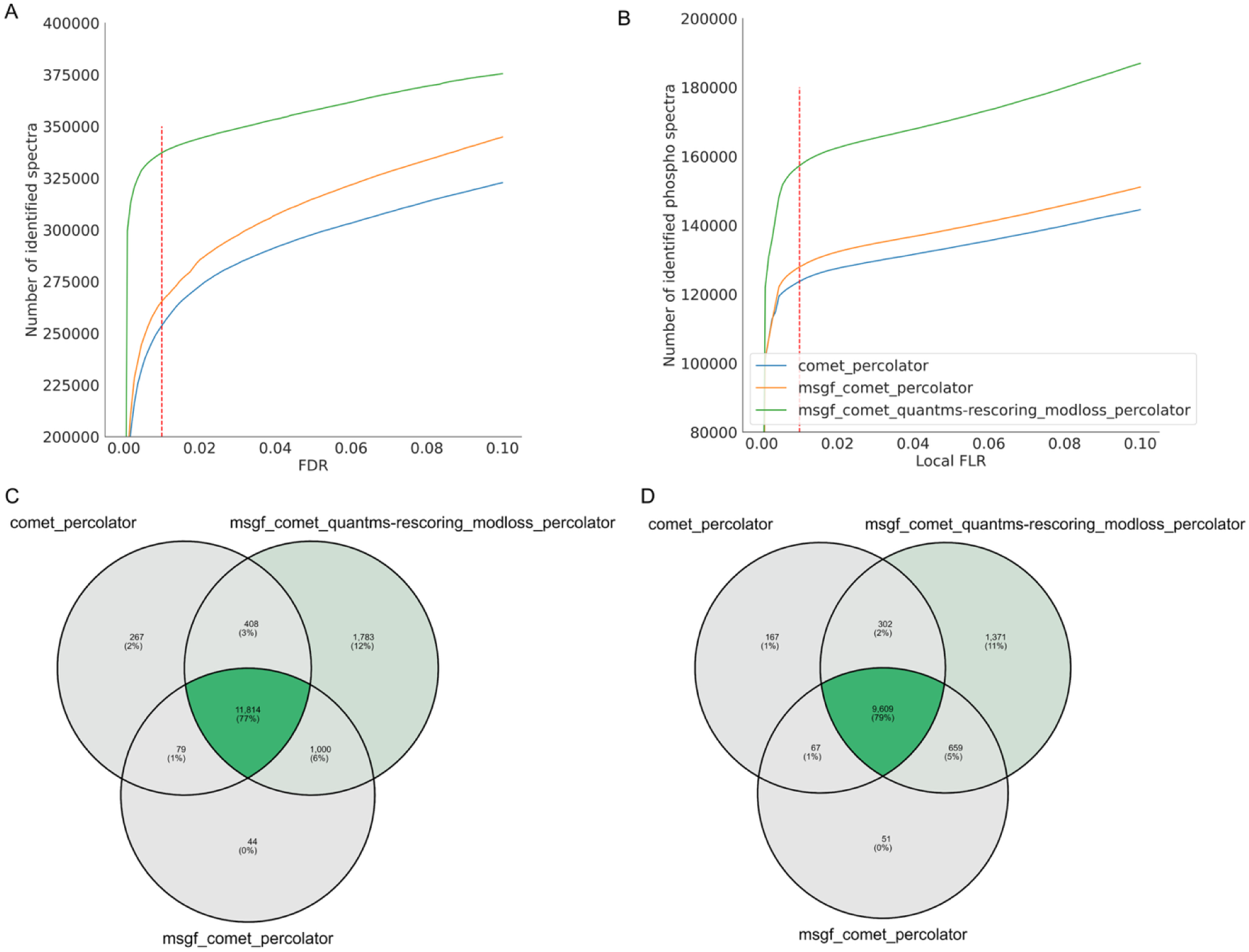
Comparison of phosphorylated peptide identification results for different workflow settings on PXD026824. (A) Number of spectra identified at different PSM FDR levels. (B) Number of phospho-PSMs at different local FLR levels; PSM filtered at 1% FDR. (C) Venn diagram of phosphorylated peptides quantified for the three settings. (D) Venn diagram of protein phosphosites at protein FDR 0.01 and FLR 0.01 for the three settings.

Because accurate phosphorylation site localization depends on robust FLR control, we also assessed how different workflows impact identifications at different FLRs (**Figure 5B**). At PSM FDR 0.01 and local FLR 0.01, the consensus workflow with quantms-rescoring features identified 20-23% more phosphorylated peptides than using a single search engine without quantms-rescoring. Under combined thresholds of PSM FDR 0.01, FLR 0.01, and protein FDR 0.01, 1,809 phosphorylated peptides were uniquely quantified in the quantms-rescoring enabled workflow; when incorporating the *modloss* feature, this number increased to 2,191 (**Figure 5C**). At the phosphosite level, the quantms-rescoring workflow uncovered 842 novel protein phosphorylation sites not detected in the other workflow configuration (**Figure 5D**), and an additional 495 sites when considering the *modloss* ions. Together, these results underscore the value of quantms-rescoring and multi-engine search for both identification site localization in phosphoproteomics, establishing quantms as a robust solution for large-scale PTM data analysis.

In addition, we also evaluated and compared FDR and FLR control for different workflow configurations on the PXD009449 synthetic (34) phosphorylated peptide dataset. The results are shown in **Supplementary Table 3**. The estimated FDR and FLR (0.07% at the PSM level and 0.4%) are larger than the actual FDP and FLR when the FDR and FLR threshold is set to 1%, and the introduction of multiple search engines and rescoring also does not disrupt the control of FDP. Importantly, combining MS-GF+ and Comet with quantms-rescoring yielded 6.5% more identified phosphorylated PSMs than using Comet alone.

### Performance of the quantms-rescoring workflow in unseen malonylation experiment

Given that the training datasets in the existing MS2 prediction models only cover common modifications, the fragmentation patterns of modified peptides and unmodified peptides exhibit varying degrees of divergence for different modifications. For example, lysine malonylation is a new important post-translational modification (PTM), which has been reported in several prokaryotic and eukaryotic species. The ProteomeTools project demonstrates that malonylated peptides drastically change the overall appearance of the spectra (median spectrum contrast angle 0.25 compared to unmodified peptides) (34). Here, we further evaluated the performance of the quantms with quantms-rescoring workflow by reanalyzing a malonylation experiment (PXD015809). The AlphaPeptDeep model is used and is fine-tuned based on ProteomeTools synthesized malonylated peptides when enabling fine-tuning. Four workflow configurations, all using Percolator in rescoring, were benchmarked: (1) Comet alone, (2) Comet and MSGF, (3) Comet and MSGF, with quantms-rescoring features, and (4) Comet and MSGF with quantms-rescoring features based on a fine-tuning model. MaxQuant is selected as a reference.

As shown in **Figure 6**, combining multiple search engines improved 12.4% PSM identifications from malonylated peptides relative to the Comet-only workflow at PSM FDR 0.01 and FLR 0.01 (**Figure 6A**). Incorporating quantms-rescoring features into the two-search-engine consensus further increased the number of identified spectra and modification sites by 25.6% and 7%, respectively. When enabling transfer learning, an additional 31.1% of PSM identifications and 21.9% modification sites from lysine malonylated peptides were achieved. Compared to MaxQuant, quantms with quantms-rescoring fine-tuning reported more 62.7% malonylation PSMs and 150% modification sites (**Figure 6B**). Then, we explored the distribution of similarity between predicted MS2 intensity and observed MS2 intensity. By fine-tuning the model to learn MS2 intensity variations of modified peptides, the cosine similarity between predicted and experimental spectra for final identification at FDR 0.01 and FLR 0.01 was enhanced from 94.9% to 98.1% (**Figure 6C**). We also observed that PSMs reported solely in configurations employing fine-tuning (0.968) exhibited greater similarity than those reported exclusively in configurations utilizing pre-trained models (0.937). The retention time predictions continue to exhibit the same trend, with all jointly identified PSMs showing RT median errors within ten minutes, and the PSMs reported solely in configurations employing fine-tuning exhibited smaller RT errors than those reported exclusively in configurations utilizing pre-trained models (**Figure 6D**).

**Figure 6:**
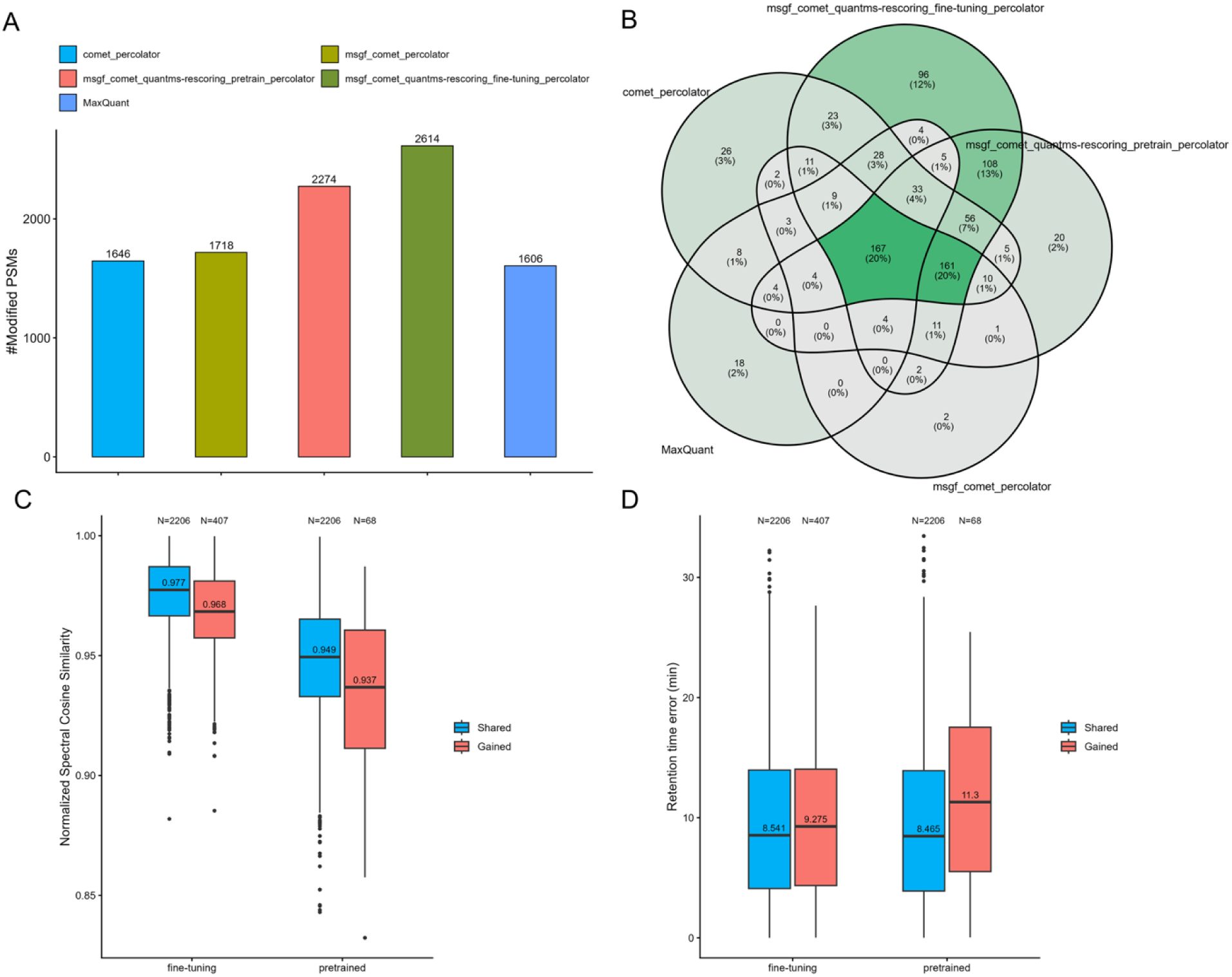
Comparison of malonylation peptide identification results for different workflow settings on PXD015809. (A) Number of modified spectra identified for five quantms configurations and MaxQuant. (B) Venn diagram of protein malonylation sites at PSM FDR 0.01, protein FDR 0.01, and FLR 0.01 for the different workflows. (C) The distribution of cosine similarity between predicted and observed modified spectra at PSM FDR 0.01 and FLR 0.01. The blue boxes represent PSMs jointly identified by the pre-trained model and the fine-tuned model. The red boxes represent the respective uniquely identified PSMs. (D) The distribution of retention time error between predicted and observed retention time for modified peptides.

We observed a similar distribution of feature weights to the results of the immunopeptidomics dataset (**Supplementary Figure 9**). It reveals strong contributions of MSGF+ features such as SpecEValue and suggests that there are limited gains when MSGF+ is used alone. In contrast, eleven AlphaPeptDeep-derived features and two DeepLC-derived features dominated the weight distribution, including *RtDiffBest* and *SpecSpearman* when Comet was used. This further demonstrates that consensus rescoring effectively exploits complementary results across different search engines.

## Discussion

Advancements in ML and deep learning-based methods in proteomics have greatly improved PSM rescoring and peptide identification (14, 22–25). As a result, several identification and quantification tools have begun integrating these models directly into their analysis workflows, including MHCQuant (25), FragPipe (24), or Mascot (42). FragPipe, for example, has introduced a new deep learning–based PSM rescoring module, MSBooster, as part of its computational platform. quantms remains one of the only open-source, community-supported workflows for quantitative proteomics and phospho-proteomics analysis in the field, enabling large-scale reanalysis of public proteomics data (8). One of the main features of quantms is that it supports open-source search engines, including MSGF+, SAGE, and Comet, and the combination of them with Percolator. The multi-search engine approaches have proved to be crucial in deep proteome studies, including proteogenomics (43, 44), and PTM analyses (45).

In this study, we developed and integrated quantms-rescoring into the quantms pipeline, an open-source cloud-based workflow for proteomics data analysis. quantms-rescoring is a lightweight module that leverages multiple machine-learning tools to seamlessly incorporate predicted ML-derived features into the quantms workflow. While conceptually related to MS²Rescore, quantms-rescoring differs in several important aspects. First, it focuses exclusively on generating PSM-level features and does not embed Percolator (19) or Mokapot (46), allowing the quantms pipeline to flexibly adopt external versions of these tools for downstream rescoring. Second, beyond the MS^2^PIP support provided by MS²Rescore, quantms-rescoring implements additional fragmentation-ion prediction capabilities through AlphaPeptDeep. AlphaPeptDeep, apart from adding more modules to the library, is technically important because, for reanalysis and also user-based analysis, users may want to perform the analysis using fragment mass errors in ppm, and MS^2^PIP only supports Daltons (Da). More importantly, as pre-trained models only cover a limited range of experimental data types, such as common PTMs, quantms-rescoring seamlessly supports model fine-tuning to generate project-specific model weights. The pre-trained models and fine-tuned models are compared on an independent test set to avoid potential fine-tuning errors. To handle multiple models and algorithms, the library provides model selection algorithms that help users of quantms to run the algorithm with the model that fits their data better. Finally, it includes extended data-cleaning procedures for search engines such as MSGF+ and introduces new feature types, including spectrum-quality metrics like signal-to-noise ratio (SNR) (36), thereby expanding the scope and adaptability of the quantms feature-generation framework (**Supplementary Table 4**). Moreover, all models are implemented to run on CPUs, which is advantageous for high-performance computing (HPC) and cloud infrastructures where GPUs are unavailable.

We evaluated its impact across multiple experimental contexts, including label-free quantification, TMT-based proteomics, immunopeptidomics, and phosphoproteomics. Our analyses show that combining multi-engine database search with both handcrafted features and features derived from retention-time and fragment-ion intensity prediction models consistently improves spectra identification and protein quantification. In particular, predicted peptide features improve Percolator’s ability to discriminate between true and false PSMs during rescoring. The resulting increase in identification confidence propagates downstream to PTM localization and quantification, leading to higher numbers of proteins with stable abundance estimates, more differentially expressed proteins, and additional phosphorylation sites. In label-free experiments, quantms-rescoring achieved 28.6% significant increases in PSM and peptide identification rates while maintaining or improving quantification reproducibility. Yang et. al. (24) observed a similar increase of 31.4% in the number of spectra identified on their LFQ immunopeptide test dataset when integrating MSBooster into FragPipe. In the TMT-labelled dataset, multiple search engines with quantms-rescoring resulted in 1369 additional quantified proteins. and revealed potentially relevant biological differences, such as novel 76 differentially expressed proteins. Similarly, in immunopeptidomics and phosphoproteomics, rescoring yielded consistently an additional 11,688 (13.6%) HLA-II potential binders and 1337 phosphosites. Similarly, the authors of MHCquant2 achieved a similar improvement (13%) in HLA-II binder identification, and it only used a search engine combined with MS²Rescore (25). This may indicate that the identification of HLA peptides can further benefit from multiple search engines’ consensus scoring. We also observed that Comet, combined with quantms-rescoring, outperformed the Comet and MS-GF+ workflow in both immunopeptidomics and phosphoproteomics analyses, yielding an 18% increase in peptide identifications and a 15% increase in PSM identifications (**Supplementary Figure 6, 7**), and quantms-rescoring reduced time by 27.6% compared to the latter (**Supplementary Table 5**). For unseen malonylation data, multiple search engines combined with quantms-rescoring fine-tuning identified 21.9% and 150% more modification sites compared to the quantms-rescoring pre-trained model and MaxQuant, respectively. Runtime benchmark indicates a modest 15%-50% increase in workflow with quantms-rescoring run time, which can be mitigated by increasing the number of quantms-rescoring parallel processes (**Supplementary Table 5**). This trade-off remains minor compared to substantial gains. quantms with quantms-rescoring provides researchers with access to multiple search engines combined with state-of-the-art models within a unified, reproducible, transparent, and scalable pipeline. This integration eliminates the need for complex manual configuration, making advanced deep learning / machine-learning tools more accessible in the context of large-scale deep proteome reanalysis. Looking ahead, extending quantms-rescoring to support a broader range of PTMs and additional DL methods -such as Prosit or a more generic approach like Koina (47) - will broaden its applicability.

## Acknowledgments

We thank all those who supported this research, including funding bodies and the proteomics community, for making proteomics data sets publicly available. Y.P-R would like to acknowledge funding from EMBL core funding, Wellcome grants (208391/Z/17/Z, 223745/Z/21/Z), and the BBSRC grant ‘DIA-Exchange’ [BB/X001911/1]. R.G., L.M., and R.B. acknowledge funding from [12B7123N,G010023N,G028821N,12A6L24N]. L.M. acknowledges funding from the Horizon Europe Projects BAXERNA 2.0 [101080544] and COMBINE [101191739], and from the Ghent University Concerted Research Action [BOF21/GOA/033]. L.M. is further supported by the CHIST-ERA project ODEEP-EU [G0GDV23N]. T.S. and O.K. acknowledge funding by the Federal Ministry of Education and Research in the frame of de.NBI/ELIXIR-DE (W-de.NBI-022). O.K. and T.S. acknowledge support by the Ministry of Science, Research and Arts Baden-Württemberg (LIBIS).

## Data Availability

The code for quantms-rescoring is available on GitHub: https://github.com/bigbio/quantms-rescoring, and the code for the quantms workflow is available on GitHub: https://github.com/bigbio/quantms. All MS/MS datasets used in this study can be found at the PRIDE partner repository or at the CPTAC repository with the following accession codes: LFQ spiked-in dataset PXD001819, HLA peptidome PXD019643, synthetic HLA-I and HLA-II peptide dataset PXD021013, TMT dataset PDC000127, synthetic phosphorylated peptide dataset PXD009449, phosphorylation dataset PXD026824, malonylation dataset PXD015809, and synthetic malonylation dataset PXD009449. quantms-rescoring version 0.0.14 is available in quantms since version v1.7 (https://github.com/bigbio/quantms/releases/tag/1.7.0)/.

## CRediT authorship contribution statement

C.D., T.S., and Y.P-R. developed all algorithms and integrated them into quantms. H.W. assisted with the quantms integration. R.G., R.B., and J.S. contributed to the integration of DeepLC, MS²PIP, and MS2Rescore into quantms-rescoring. A.L., C.D., and Y.P-R. designed the benchmarking strategy and curated several datasets. T.S., C.D., and Y.P-R. wrote the original manuscript. O.K., L.M., F.H., M.B., and L.X. supported the work and provided essential resources. All authors read, contributed to, and approved the final manuscript.

