## Supplementary Notes for "quantms-rescoring enables deep proteome coverage across protein quantification, immunopeptidomics, and post-translational modifications experiments"

### **Supplemental Figure 1**: Comparison of identification and quantification results for different workflow settings on the PXD001819 UPS1 spike-in dataset.

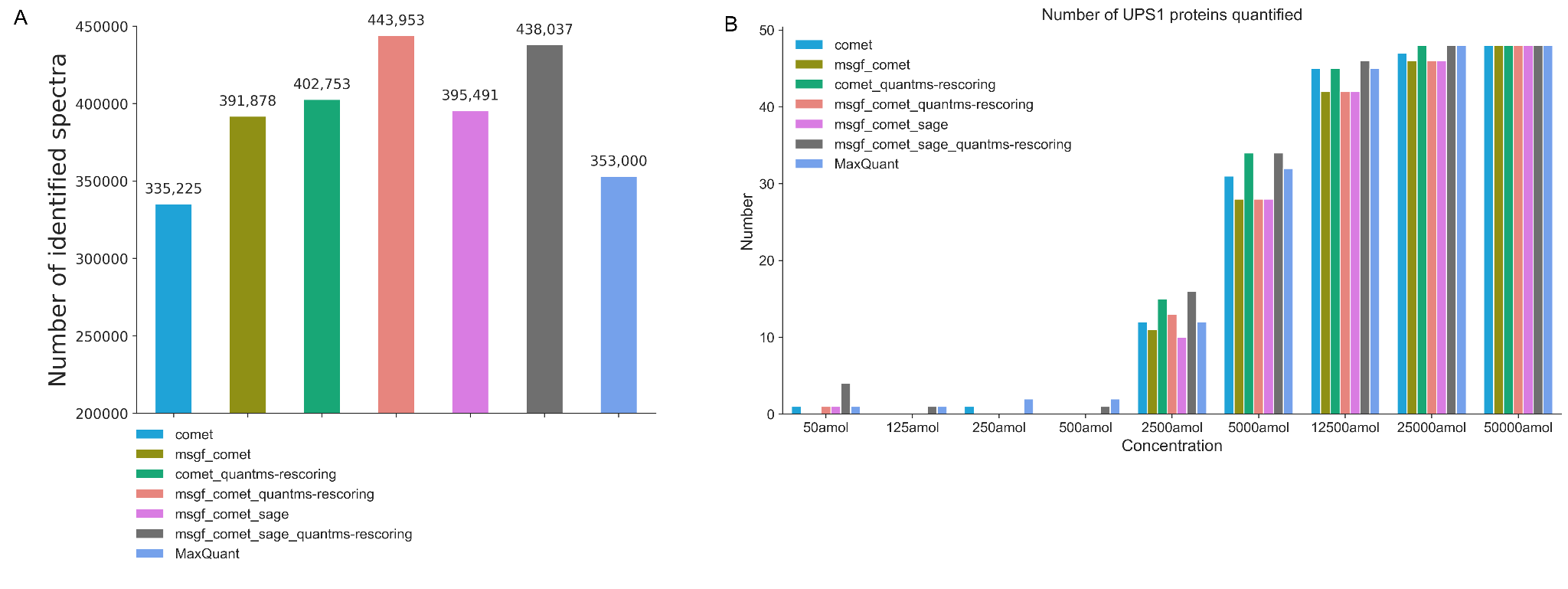

**Supplemental Figure 1**: Comparison of identification and quantification results for different workflow settings on the PXD001819 UPS1 spike-in dataset (A) The number of identified spectra at 0.01 FDR levels for different workflow settings. (B) Results for the PXD001819 dataset showing protein quantification. Six workflow configurations that employ Percolator during post-scoring were compared: (1) Comet alone, (2) two search engines, (3) Comet with quantms-rescoring features, (4) two search engines with quantms-rescoring features, (5) three search engines, and (6) three search engines with quantms-rescoring features.

### **Supplemental Figure 2**: The top 20 Percolator feature weights dataset PXD001819.

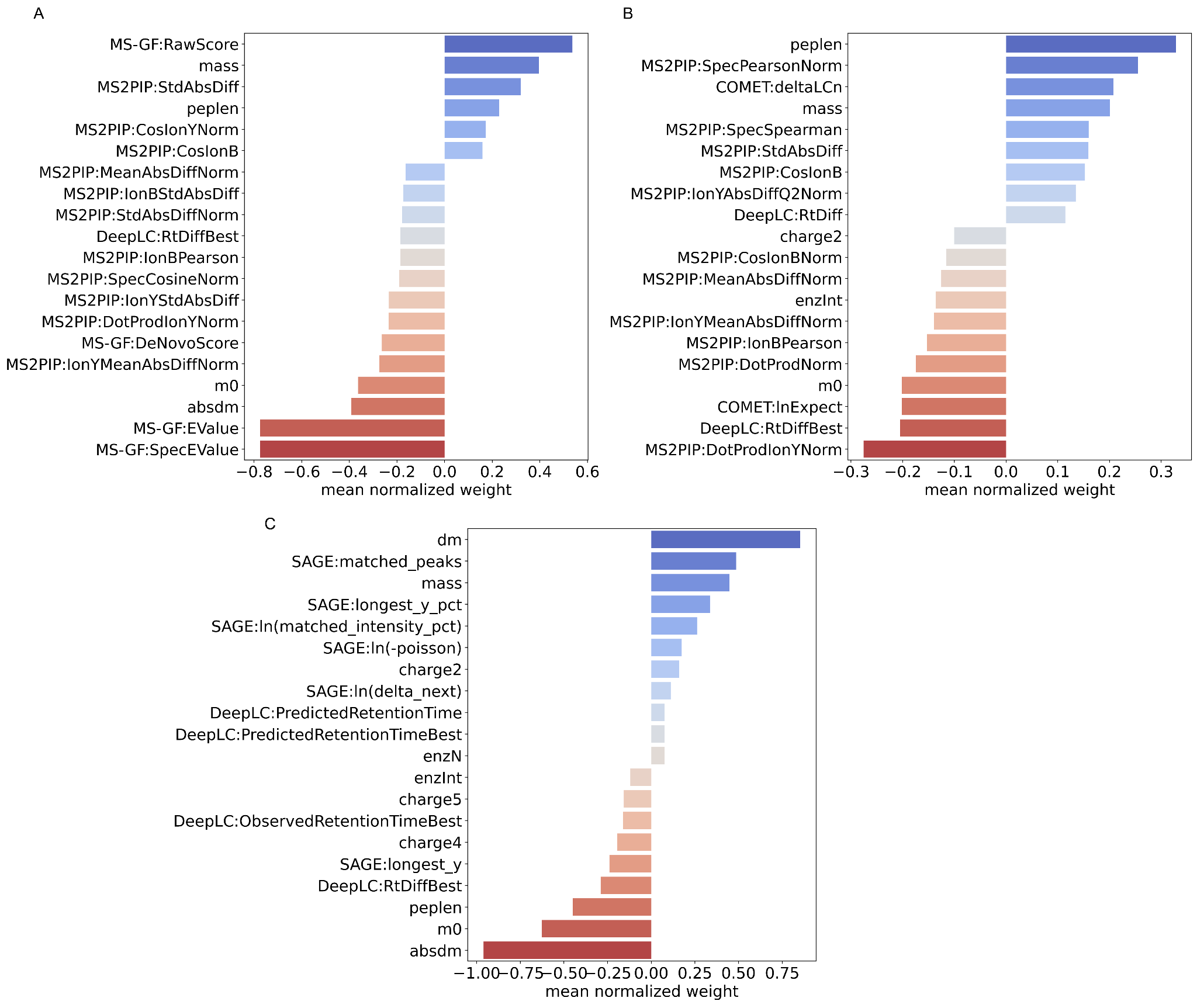

**Supplemental Figure 2**: The top 20 Percolator feature weights for MSGF+ (A), Comet (B), and SAGE (C) combined with Percolator in the quantms-rescoring workflow (dataset PXD001819). Features with weights farther from zero - either positive or negative - contribute more strongly to the final Percolator classification.

### **Supplemental Figure 3**: Comparison of identification and quantification results for different workflow settings on the CPTAC TMT dataset PDC000127.

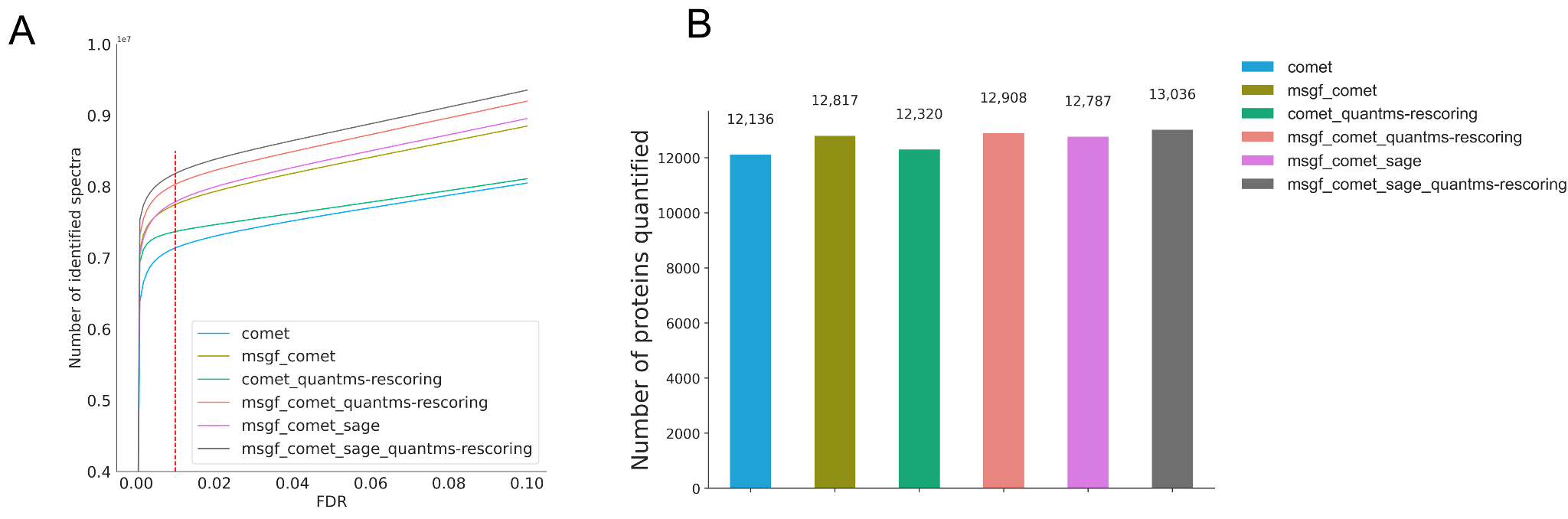

**Supplemental Figure 3**: Comparison of identification and quantification results for different workflow settings on the CPTAC TMT dataset PDC000127. (A) Number of identified spectra at different FDR thresholds and six workflow settings. (B) The barplot of quantified proteins for six workflow settings. Six workflow configurations that employ Percolator during post-scoring were compared: (1) Comet alone, (2) two search engines, (3) Comet with quantms-rescoring features, (4) two search engines with quantms-rescoring features, (5) three search engines, and (6) three search engines with quantms-rescoring features.

### **Supplemental Figure 4**: Top 20 feature weights assigned by Percolator for dataset PDC000125.

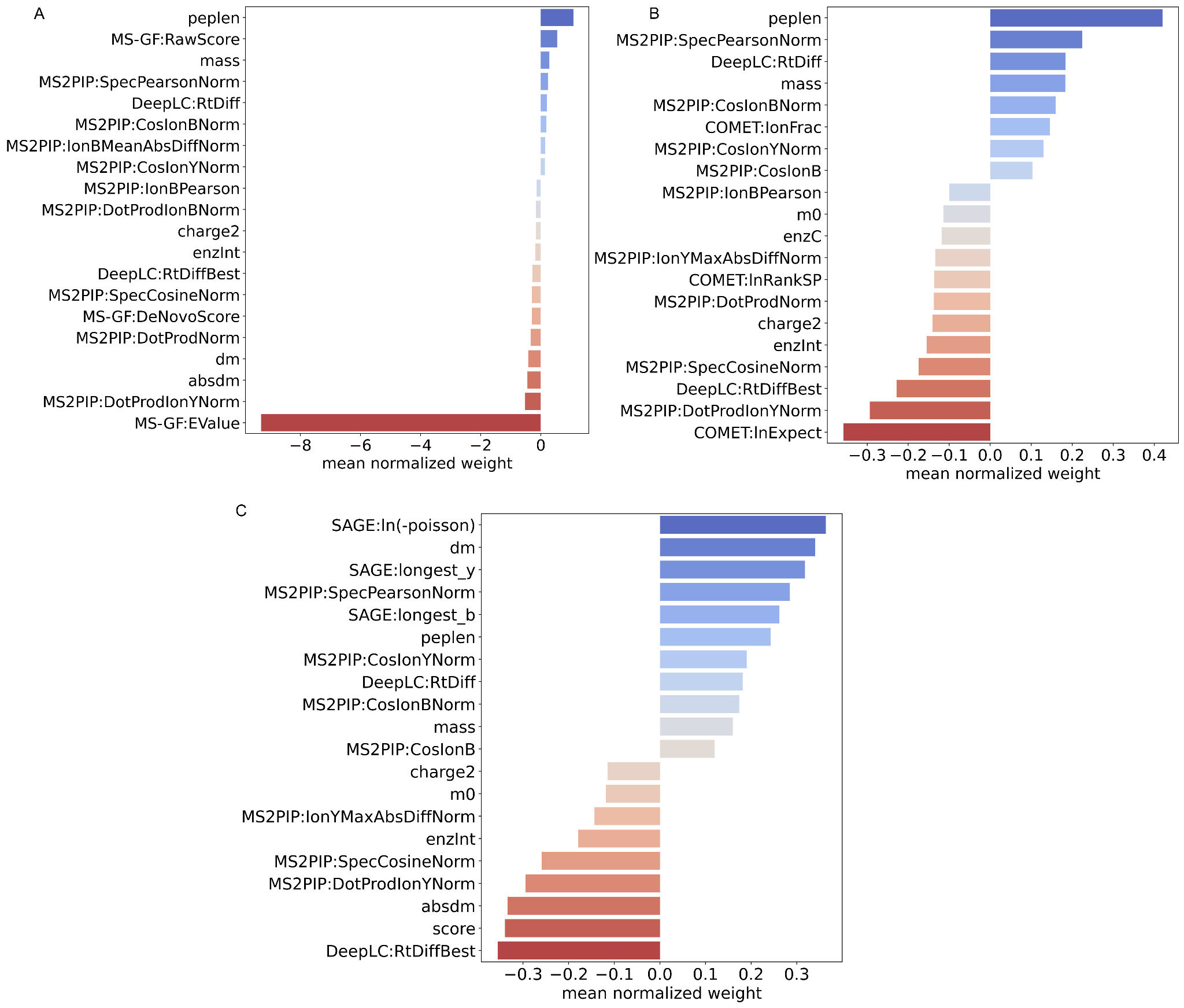

**Supplemental Figure 4**: Top 20 feature weights assigned by Percolator for MSGF+ (A), Comet (B), and SAGE (C) in the quantms-rescoring workflow applied to CPTAC PDC000125 TMT datasets.

### **Supplemental Table 1**: The added spectra feature by quantms-rescoring.

| **Feature Name** | **Definition** |
| --- | --- |
| **signal to noise (SNR)** | The signal-to-noise ratio calculated as the maximum intensity divided by the root mean square deviation of all intensities for a spectrum. |
| **spectral entropy** | All peak intensities are normalized by total ion current, and the entropy of all normalized peak intensities in the spectrum is calculated. |
| **fraction tic top 10** | Fraction of total ion current covered by the top 10 peaks. |
| **weighted std mz** | Compute the weighted mean m/z by summing each mass-to-charge value multiplied by its normalized intensity. Then calculate the weighted standard deviation by subtracting the weighted mean from each m/z, squaring, multiplying by normalized intensity, summing, and taking the square root. |

### **Supplemental Figure 5**: Top 20 normalized feature weights assigned by Percolator for dataset PXD019643.

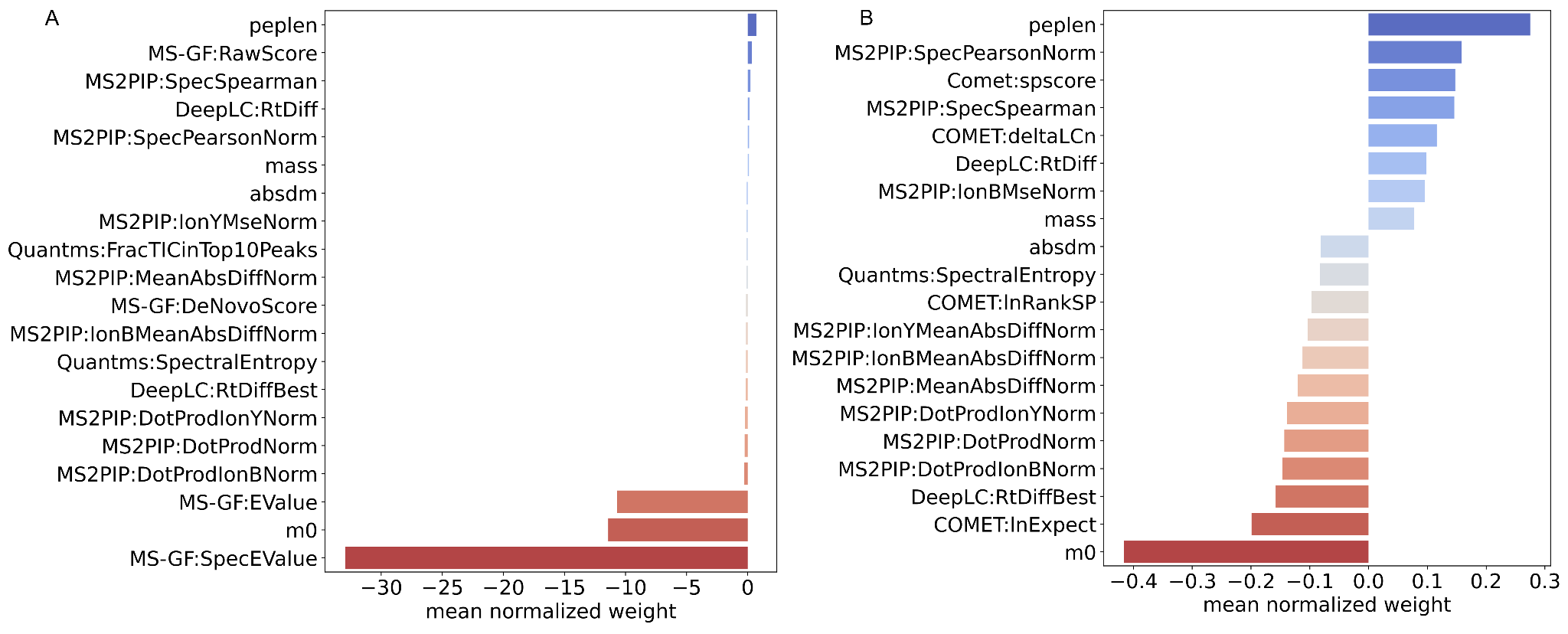

**Supplemental Figure 5**: Top 20 normalized feature weights assigned by Percolator for MSGF+ (A) and Comet (B) identification results in the PXD019643 HLA Class II immunopeptidomics dataset.

**Supplemental Figure 6**: Comparison of HLA-II immunopeptide identifications for five quantms workflow configurations on PXD019643.

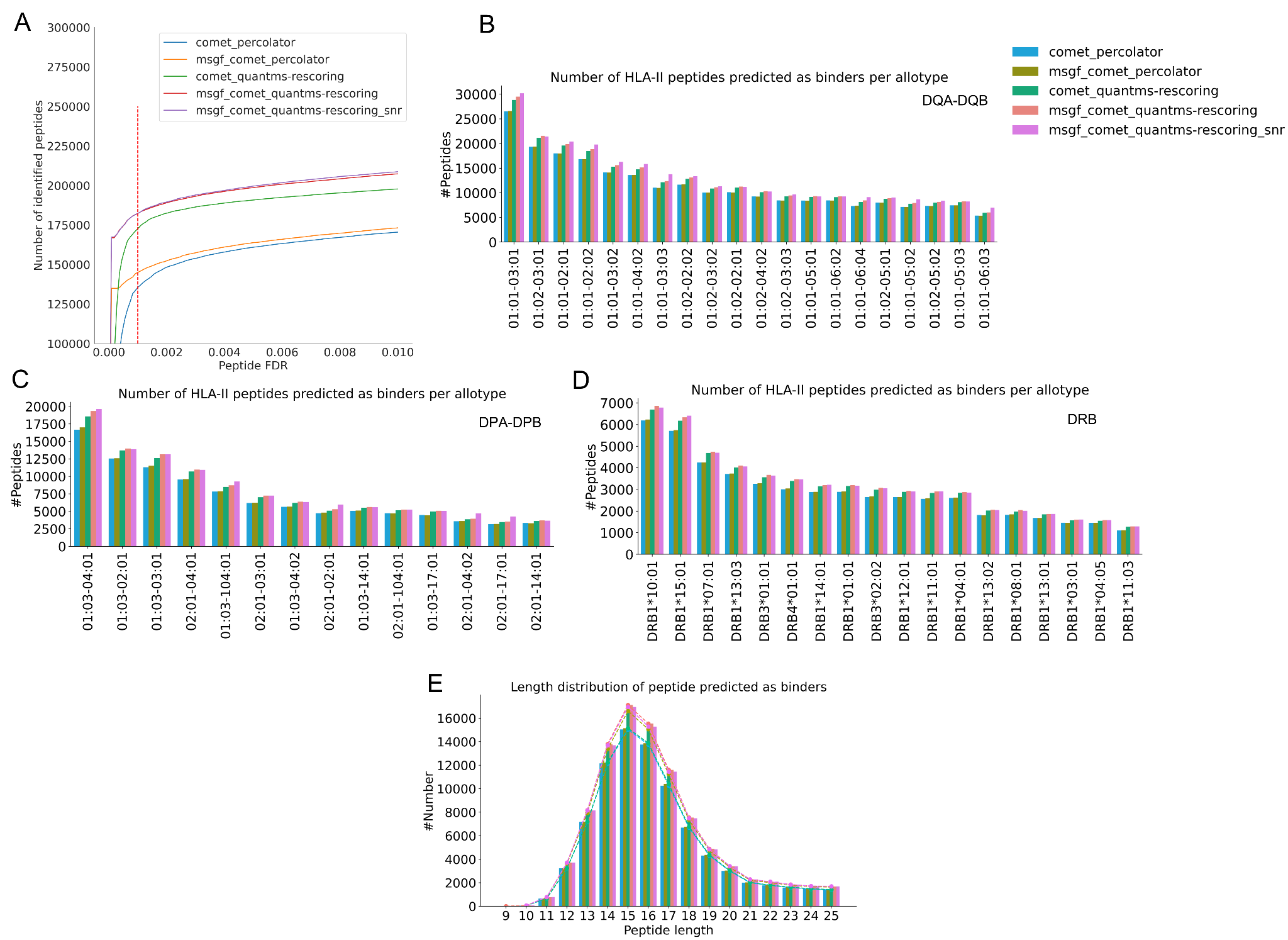

**Supplemental Figure 6**: Comparison of HLA-II immunopeptide identifications for five quantms workflow configurations on PXD019643. (A) Peptide identifications across FDR thresholds. (B–D) Global overview of HLA-II predicted binders distributed across HLA molecules. HLA binding prediction was performed with NetMHCIIpan-4.3. HLA class II peptides were defined as binders when the percentile rank is less than or equal to 5. (E) Length distribution of identified HLA-II peptides predicted as binders from all samples was analyzed.

### **Supplemental Table 2**: Comparison of false discovery proportion (FDP) control at the PSM and peptide levels for the PXD021013 synthetic HLA peptide dataset.

| **Workflow Configuration** |  | **Total identified Spectra** | **False Spectra Identification** | **FDP at PSM level** |
| --- | --- | --- | --- | --- |
| msgf_comet_quantms-rescoring_snr | Estimated PSM FDR 1% | 593052 | 7527 | 1.2% |
|  |  | **Total identified peptide** | **False identified peptide** | **FDP at peptide level** |
|  | Estimated peptide FDR 1% | 37114 | 631 | 1.7% |
|  |  | **Total identified Spectra** | **False Spectra Identification** | **FDP at PSM level** |
| msgf_comet | Estimated PSM FDR 1% | 497433 | 6214 | 1.2% |
|  |  | **Total identified peptide** | **False identified peptide** | **FDP at peptide level** |
|  | Estimated peptide FDR 1% | 31856 | 530 | 1.6% |
|  |  | **Total identified Spectra** | **False Spectra Identification** | **FDP at PSM level** |
| comet | Estimated PSM FDR 1% | 500688 | 6394 | 1.2% |
|  |  | **Total identified peptide** | **False identified peptide** | **FDP at peptide level** |
|  | Estimated peptide FDR 1% | 32331 | 532 | 1.6% |

**Supplemental Table 2**: Comparison of false discovery proportion (FDP) control at the PSM and peptide levels for the PXD021013 synthetic HLA peptide dataset. Three workflow configurations that employ Percolator during post-scoring were compared. 1% FDR are applied at the PSM level and peptide level.

### **Supplemental Table 3**: The results of FDR quality control at the PSM level and of FLR control at the site localization level on the PXD009449 synthetic phosphorylated peptides datasets.

| **Workflow configuration** |  | **Total Phospho Spectra** | **False Phospho Spectra identification** | **FDP at PSM level** |
| --- | --- | --- | --- | --- |
| comet_msgf+_quantms-rescoring | Estimated PSM FDR 1% | 102974 | 84 | 0.07% |
|  |  | **Total Phospho Spectra** | **False Phospho sites localization** | **Real FLR** |
|  | Estimated global FLR 1% | 86053 | 346 | 0.40% |
|  |  | **Total Phospho Spectra** | **False Phospho Spectra identification** | **FDP at PSM level** |
| comet_msgf+ | Estimated FDR 1% | 102792 | 74 | 0.07% |
|  |  | **Total Phospho Spectra** | **False Phospho sites localization** | **Real FLR** |
|  | Estimated global FLR 1% | 86029 | 341 | 0.39% |
|  |  | **Total Phospho Spectra** | **False Phospho Spectra identification** | **FDP at PSM level** |
| comet | Estimated PSM FDR 1% | 95679 | 15 | 0.01% |
|  |  | **Total Phospho Spectra** | **False Phospho sites localization** | **Real FLR** |
|  | Estimated global FLR 1% | 80766 | 297 | 0.36% |

**Supplemental Table 3**: The results of FDR quality control at the PSM level and of FLR control at the site localization level on the PXD009449 synthetic phosphorylated peptides datasets. Three workflow configurations that employ Percolator during post-scoring were compared. 1% FDR is applied at the PSM level, and 1% FLR is applied at the localization level.

### **Supplemental Figure 7**: Comparison of phosphorylated peptide identification results for different workflow settings on PXD026824.

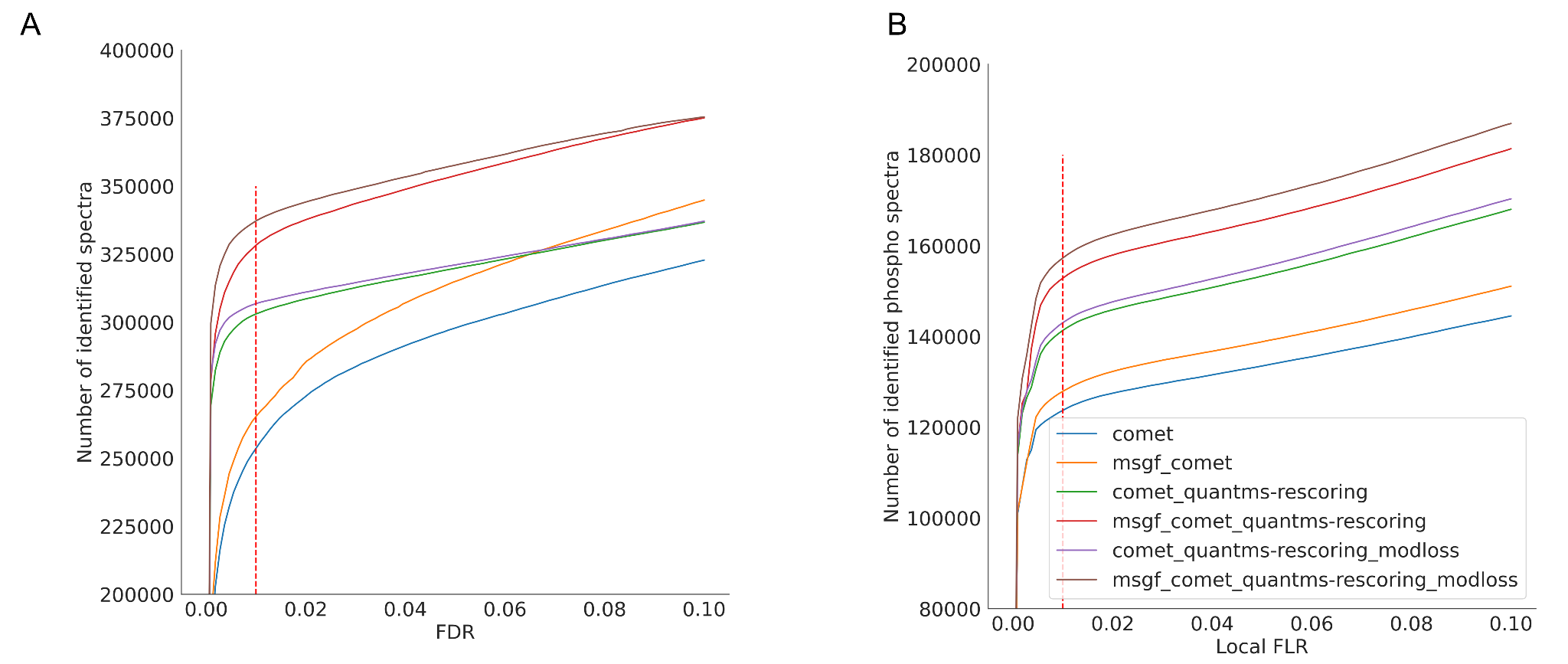

**Supplemental Figure 7**: Comparison of phosphorylated peptide identification results for different workflow settings on PXD026824. (A) Number of spectra identified at different PSM FDR levels. (B) Number of phospho-PSMs at different local false localization rate levels. Six workflow configurations that employ Percolator during post-scoring were compared. 1% FDR is applied at the PSM level, and 1% FLR is applied at the localization level.

### **Supplemental Figure 8**: Top 20 feature weights assigned by Percolator dataset PXD026824.

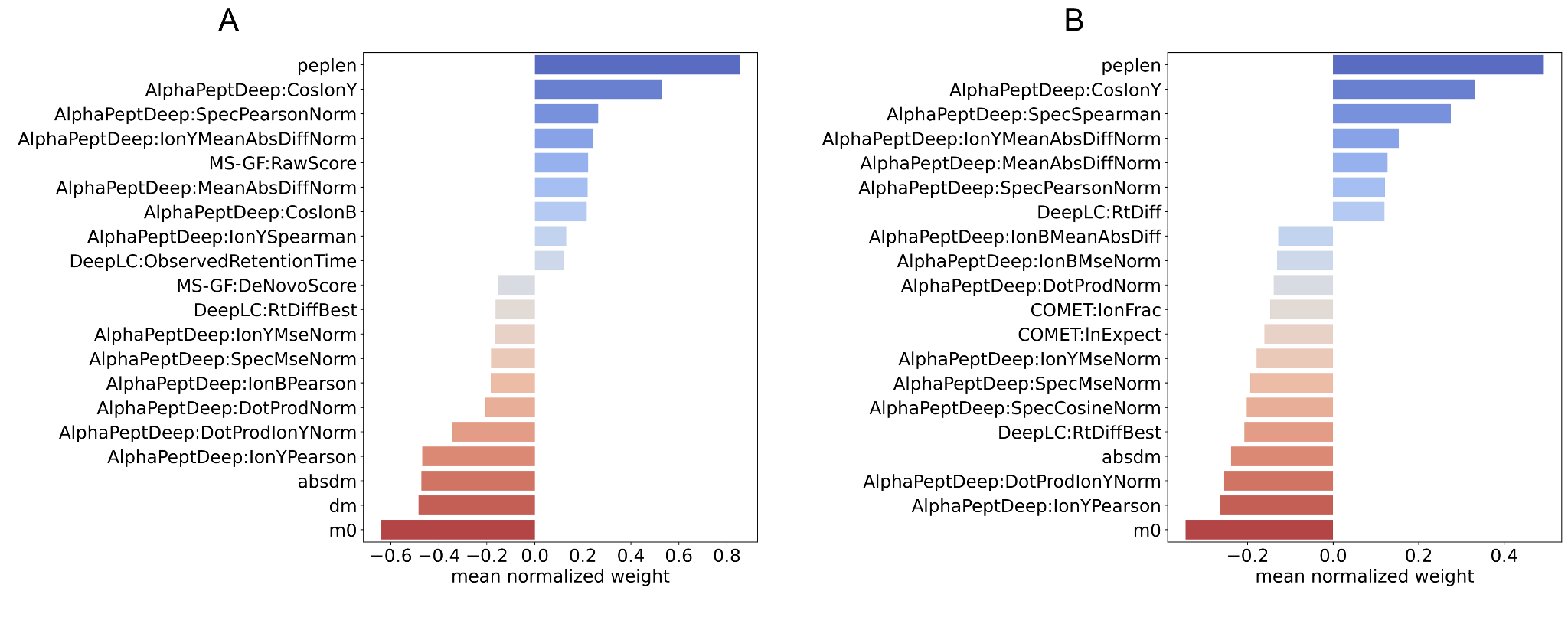

**Supplemental Figure 8**: Top 20 feature weights assigned by Percolator for the MSGF+ (A) and Comet (B) search engine feature vectors, using settings optimized for the post-translational modification dataset (PXD026824).

### **Supplemental Table 4**: The characteristics of quantms-rescoring, MS²Rescore and MSBooster.

|  | **quantms-rescoring** | **MS²Rescore** | **MSBooster** |
| --- | --- | --- | --- |
| **Language** | Python | Python | Java |
| **Multi ML-based method (if more than one is supported)** | ☑ | ☑ | ☑ |
| **model selection optimization** | ☑ | ☒ | ☑ |
| **validation safeguard step** | ☑ | ☒ | ☒ |
| **Standalone Container ready** | ☑ | ☑ | ☒ |
| **Support input file** | idXML, mzML | More than 20 file types | .pin,  mzML/MGF |
| **Include Percolator or Mokapot** | ☒ | ☑ | ☒ |
| **Include additional spectrum features** | ☑ | ☒ | ☒ |
| **feature filter capabilities** | ☑ | ☑ | ☒ |
| **Supported mass tolerances units** | ppm/Da | ppm | ppm/Da |

### **Supplemental Table 5**: Total runtime, maximum memory, and maximum CPU consumption in different configurations.

| Datasets | Configuration | Runtime | Maximum Memory | Maximum CPU consumption |
| --- | --- | --- | --- | --- |
| PDC000127 | comet_percolator | 5h 10m | 36 GB | 7 |
| PDC000127 | comet_msgf_sage_percolator | 6h 28m | 37 GB | 7 |
| PDC000127 | comet_msgf_sage_quantms-rescoring_percolator | 9h 39m | 37 GB | 8 |
| PXD019643 | comet_percolator | 2h 05m | 7.7 GB | 6 |
| PXD019643 | comet_msgf_percolator | 4h 34m | 7.7 GB | 8 |
| PXD019643 | comet_msgf_quantms-rescoring_percolator | 5h 08m | 40.8 GB | 8 |
| PXD026824 | comet_percolator | 24m | 4.8 GB | 5 |
| PXD026824 | comet_msgf_percolator | 38m | 7.7 GB | 6 |
| PXD026824 | comet_msgf_quantms-rescoring_percolator | 55m | 68.3 GB | 6 |

**Supplemental Table 5**: Total runtime, maximum memory, and maximum CPU consumption in different configurations. The submitted job queue of quantms was limited to 64. The memory and CPU values represent the maximum resources required for a single job.
